# The CXCL10/CXCR3 axis is essential for sustaining immunological dormancy in breast cancer

**DOI:** 10.1101/2025.09.26.678769

**Authors:** Alev Yilmaz, Lisa Haerri, Mateo Estrella Granda, Oriana Coquoz, Qiang Lan, Curzio Rüegg

## Abstract

Immune surveillance plays a pivotal role in controlling tumor emergence, dormancy and progression, including in breast cancer. Despite its potential clinical relevance, the mechanisms governing dormancy initiation, maintenance and escape, as well as the molecular mediators involved, remain poorly understood. Here, we identify the interferon-inducible chemokine CXCL10 and its receptor CXCR3 as key regulators of immunological dormancy in breast cancer. By transcriptomic profiling we observed high expression of *Cxcl10* in dormant cells in two different orthotopic, syngeneic models of breast cancer dormancy (D2.0R and 4T1-MR20). Genetic silencing of *Cxcl10* in dormant cells or pharmacological blockade of CXCR3 *in vivo*, led to early tumor onset and rapid growth in immunocompetent mice. In contrast dormant cells effectively formed tumors in immune-deficient mice independently of *Cxcl10* status, demonstrating that the CXCL10/CXCR3 axis-mediated dormancy requires a functional immune system. Further analysis confirmed that *Cxcl10* silencing altered the local immune microenvironment, reducing CD4^+^ and CD8^+^ T cell infiltration while increasing the presence of granulocytic Myeloid Derived Suppressor Cells and Natural Killer cells. Moreover, *Cxcl10* silencing significantly increased the burden of tumor cells disseminated to the lung. Leveraging on these findings, we identified a CXCL10-mediated dormancy signature that predicts improved overall survival in triple-negative breast cancer (TNBC) patients. Our findings have identified a new mechanism modulating breast cancer dormancy with important clinical implications: the CXCL10/CXCR3 axis as a potential therapeutic target for improving survival of patients with TNBC, and the CXCL10-dependent dormancy signature as a tool for identifying these patients.

## Introduction

Despite significant advances in early detection and treatment, breast cancer remains the leading cause of cancer-related mortality among women worldwide [1, 2]. A major contributor to this dismal clinical situation is the ability of disseminated tumor cells (DTCs) to persist in a dormant state for years or even decades before reactivating to form overt metastases that will eventually kill the patient [3–5]. Tumor dormancy poses significant barriers to long-term therapeutic success, as dormant tumor cells are often beyond the detection resolution of standard diagnostic tools and are largely resistant to therapies targeting proliferating cells [5–8]. The biological underpinnings of tumor dormancy, particularly in the context of breast cancer, have therefore become a critical focus of cancer research recently.

Tumor dormancy is categorized into three types: cellular dormancy (cell-intrinsic proliferative arrest), angiogenic dormancy (growth limitation due to vascular insufficiency), and immunological dormancy, where immune surveillance constrains tumor outgrowth [3, 9–11]. While cellular dormancy has been extensively studied using *in vitro* and *in vivo* models, immunological dormancy, where cytotoxic lymphocytes and innate effectors mediate a dynamic equilibrium that restrain the overall growth of tumor mass, has only recently gained wider attention [10–12].

Early support for immunological dormancy came from observations in immunocompetent murine models, where dormant tumors re-emerged following immunosuppressive treatments or when grafted in immune-deficient hosts [13, 14]. More recently, transcriptomic and functional studies have implicated type I interferon (IFN) signaling as a critical pathway mediating dormancy-associated immune responses [15–18]. Interferon-regulatory factor 7 (IRF7) and its downstream effector IFN-β have been shown to maintain dormancy in breast cancer models by enhancing immune-mediated tumor suppression [19, 20]. Yet, the downstream mediators executing these effects remain uncharacterized.

Among the potential candidates, chemokines, intercellular signaling molecules central to immune cell trafficking and tumor-immune interactions, have emerged as critical players in anti-tumor immune response [21–24]. In particular, IFN-inducible chemokines CXCL9, CXCL10, and CXCL11, which signal via a common receptor, CXCR3, have been implicated in promoting antitumor immunity by recruiting effector T cells to sites of tumor development [25, 26]. Intriguingly, their contribution to enforcing, sustaining, or disrupting immunological tumor dormancy has remained unexplored.

Recent studies have begun to identify links between chemokine expression and therapeutic efficacy in breast cancer [25, 27–29]. Notably, patients with high CXCL9/CXCL10/CXCR3 expression in their tumors showed longer survival after pembrolizumab (an anti-PD-1 checkpoint inhibitor) treatment in triple negative breast cancer (TNBC) [30]. Yet, these associations are often observational and lack mechanistic interrogation. Adding to the complexity is the fact that chemokines such as CXCL10 can exert context-dependent effects, facilitating immune infiltration and anti-tumor immune response in some settings, but also facilitating disease progression in others by promoting tumor cell proliferation, migration and invasion [31–33]. Therefore, investigating the role of these chemokines in dormancy-specific models is required to elucidate their precise functions in tumor-immune equilibrium.

In this study, we address the role of CXCL10/CXCR3 axis in immunological breast cancer dormancy by leveraging established murine models of dormancy, transcriptomic profiling, and both genetic and pharmacological perturbations. Our results demonstrate that activation of the CXCL10/CXCR3 axis is essential for maintaining breast cancer dormancy through an effective anti-tumor immune mechanism, but is not sufficient to induce tumor dormancy in non-dormant cells. A CXCL10-mediated dormancy signature predicts improved overall survival in TNBC patients supporting the clinical relevance of these observations.

## Results

### Cxcl9, Cxcl10 and Cxcl11 chemokines are preferentially expressed in dormant breast cancer cells

To investigate the molecular mechanism underlying breast cancer immunological dormancy, we conducted transcriptomic profiling of dormant D2.0R cells and their non-dormant counterparts D2A1. While D2.0R cells were originally reported as a model of cellular dormancy [34, 35], we subsequently reported that dormant D2.0R cells elicit an effective tumor-suppressive immune response through a constitutively active IRF7/IFN-β axis implicating immunological dormancy in D2.0R dormancy *in vivo* as well [20].

Consistent with these findings, gene set enrichment analysis (GSEA) demonstrated significant upregulation of the HALLMARK G2M CHECKPOINT and EF2 TARGETS pathways in dormant D2.0R cells (Figure 1A and Supplementary Table 1), aligning with their proliferation arrest. However, INTERFERON ALPHA and INTERFERON GAMMA pathways were remarkably the most highly enriched, further confirming the role of IFN signalling in mediating immune-dependent dormancy.

**Figure 1.**
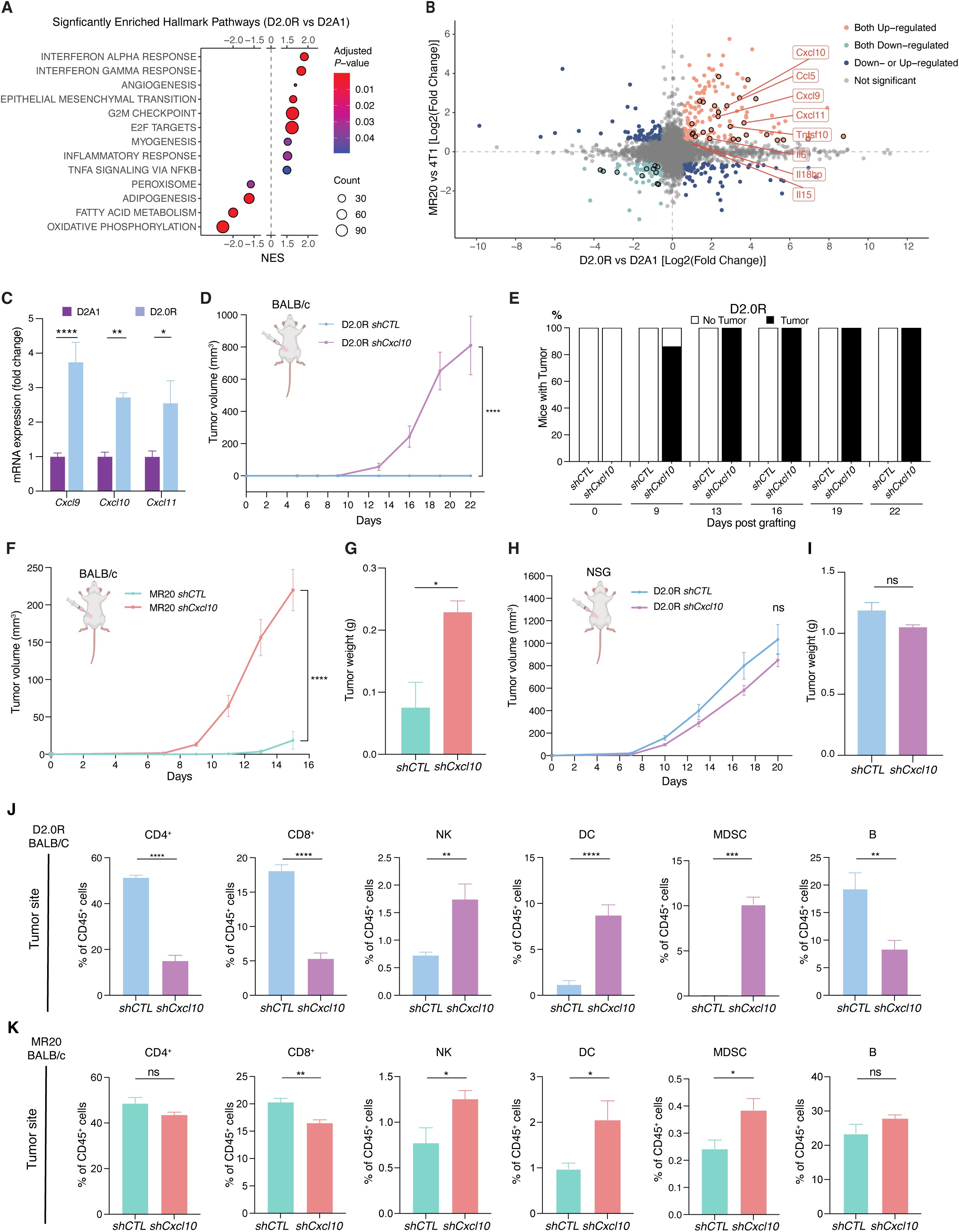
CXCL10 is essential for maintaining immunological dormancy in breast cancer by altering local immune environment. **A** Gene set enrichment analysis (GSEA) of RNAseq data from D2.0R *vs* D2A1 cells showing significantly enriched Hallmark pathways. Significance is color-coded and the number of counts indicated by the sized of the dots. **B** Dot plot showing the differentially expressed genes (DEGs) comparing D2.0R *vs* D2A1 (x-axis) and MR20 *vs* 4T1 (y-axis). The DEGs encoding ligand proteins were highlighted with black circle. The core enrichment ligand genes in INTERFERON ALPHA and GAMMA RESPONSE pathways extracted from GSEA performed in **A** are annotated with text labels (Full gene list in Supplementary Table 1). **C** Relative *Cxcl9*, *Cxcl10* and *Cxcl11* mRNA expression in D2A1 and D2.0R cell lines examined by RT-qPCR, whereby expression in D2A1 is adjusted to 1. n= 3/group. **D, E** Primary tumor growth measured by volume (**D**) and percentage of mice developing tumors over time (**E**) of D2.0R *shCTL* and *shCxcl10* cells injected into 4^th^ mammary fat pad of immune competent BALB/c mice. n= 7-8/group. **F, G** Primary tumor growth measured by volume (**H**) and tumor weight at the end of experiment (**I**) of MR20 *shCTL* and *shCxcl10* cells injected into 4^th^ mammary fat pad of immune deficient BALB/c mice. n= 8/group. **H, I** Primary tumor growth measured by volume (**F**) and tumor weight at the end of experiment (**G**) of D2.0R *shCTL* and *shCxcl10* cells injected into 4^th^ mammary fat pad of immune deficient NSG mice. n= 8/group. **J, K** Frequency of immune cells populations in the primary tumor site of BALB/c mice orthotopically injected with *shCTL* or *shCxcl10* tumor cells derived from D2.0R (**J**) and MR20 (**K**), as determined by flow cytometry analysis. Results are expressed as percentage of the indicated cell populations within CD45^+^ cells. n= 6-8/group. Data are represented as mean ± SEM. *P* values were calculated using two-way ANOVA with Sidak’s multiple-comparisons test (**D, F, H**); or unpaired two-tailed Student’s *t* test (**C, G, I, J & K**). ns, not significant; *, *P* < 0.05; **, *P* < 0.01; ***, *P* < 0.0005; ****, *P* < 0.0001.

By setting the threshold of Fold Change > 1.5, adjusted p-value < 0.05 and average expression > 50, 1443 significantly differentially expressed genes (DEGs) were identified when comparing D2.0R with D2A1 cells (Supplementary Table 1). To further identify the potential mediator of IFN-mediated immune response, we focused on the eight ligand proteins within the core enriched genes in the HALLMARK INTERFERON ALPHA and INTERFERON GAMMA pathways as identified in Figure 1A (Supplementary Table 1): *Cxcl9*, *Cxcl10*, *Cxcl11*, *Ccl5*, *Tnfsf10*, *Il6*, *Il18bp* and *Il15* (Figure 1B and Supplementary Figure 1A). Among them, *Cxcl9*, *Cxcl10*, *Cxcl11*, *Ccl5*, *Tnfsf10*, *Il6* were also significantly upregulate in MR20 (Figure 1B and Supplementary Table 1), a previously reported model of chemotherapy-induced immunological dormancy derived from 4T1 cells [20]. Notably, *Cxcl9*, *Cxcl10,* and *Cxcl11* belong to the same chemokine family that binds to a common receptor, CXCR3 [31, 36]. As *Cxcl10* emerged as the most abundant expressed ligand (Supplementary Figure 1A) and was highly upregulated in dormant (D2.0R and MR20) tumor cells comparing with non-dormant (D2A1 and 4T1) cells (Figure 1C and Supplementary Figure 1B), we decided to focus on CXCL10 for the subsequent functional analyses.

### CXCL10 is essential for maintaining immunological dormancy in vivo

To interrogate the function of CXCL10 in maintaining breast cancer dormancy, we silenced expression of *Cxcl10* in both D2.0R and MR20 cells using a shRNA-mediated knockdown (KD) approach. The KD efficiency was confirmed by qRT-PCR (Supplementary Figure 1C and D). When D2.0R *shCxcl10* tumor cells were orthotopically injected into immune competent BALB/c mice, tumor onset was observed at day 9 post-grafting and by day 13, all mice had developed palpable tumors (Figure 1D and E). In contrast, none of the mice injected with D2.0R *shCTL* developed tumors during the experimental timeframe. As observed throughout the experimental period, D2.0R *shCxcl10* tumor volume increased progressively, exhibiting similar growth kinetics as D2A1 cells in immunocompetent mice, as described in previous studies (Figure 1D) [20]. Similar findings were obtained using the chemotherapy-induced dormancy model, where all mice grafted with MR20 *shCxcl10* cells developed primary tumors shortly after grafting, while the MR20 *shCTL* group exhibited a marked delay in tumor onset as previous reported [20] (Figure 1F, G and Supplementary Figure 1E).

To corroborate the involvement of the immune system in CXCL10-mediated dormancy, we compared tumor growth of orthotopic grafted D2.0R *shCTL* and D2.0R *shCxcl10* cells in immune-compromised NOD-SCID common gamma 2 chain-deficient (NSG) mice, which lack mature T, B and NK cells and display impaired DC and macrophage functions [37]. All mice in both groups developed tumors 7 days post-grafting (Supplementary Figure 1F) and no significant difference in tumor growth was observed between D2.0R *shCTL*- and D2.0R *shCxcl10*-injected mice (Figure 1H and I).

Taken together these results indicate that CXCL10 mediates an effective anti-tumor immune response-dependent state of dormancy.

### Cxcl10 silencing suppresses T cell and B cell infiltration and promotes the accumulation of CD11b^+^Gr1^+^ myeloid cells in the tumor microenvironment

To further elucidate the mechanism underlying CXCL10-mediated immunological dormancy *in vivo*, we characterized immune cells in the tumor site and peripheral blood in the mice orthotopically injected with D2.0R *shCTL* and D2.0R *shCxcl10* cells (Figure 1J and Supplementary Figure 1G). In tumor-injected fat pad, both CD4^+^ and CD8^+^ T cells, as well as B (B220^+^) lymphocytes were significantly reduced upon suppression of *Cxcl10* (Figure 1J). Conversely, a significant increase of locally infiltrated natural killer (NK, CD49b^+^) cells, dendric cells (DCs, CD11c^+^) and Myeloid-Derived Suppressor Cells (MDSC, CD11b^+^Gr1^+^) were observed in the D2.0R *shCxcl10* tumors. A similar observation was obtained in the MR20 model, with reduced CD8^+^ T lymphocytes and increased NK cell infiltrating MR20 shCXCL10 tumors compared to controls (Figure 1K). On the other hand, only mild alterations in systemic immune response were detected in the blood of *shCxcl10* tumor-grafted mice. In the D2.0R model, this is characterized by a decrease in B cells and MDSC compared to controls (Supplementary Figure 1G). In contrast, the MR20 model exhibited a decrease in NK cells alongside an increase in DCs (Supplementary Figure 1H).

These findings indicate that downregulation of tumor-derived CXCL10 profoundly reshapes the local immune landscape, diminishing the presence of T and B lymphocytes while promoting recruitment of innate and immunosuppressive cell populations. This potent local effect, however, only partially extend to circulating immune cells.

### Reduced Cxcl10 expression in D2.0R cells increases tumor cell dissemination in the lung

To interrogate whether *Cxcl10* had an impact on metastasis, we analyzed the lungs from tumor bearing immunocompetent mice. Histological analysis revealed no evidence of macroscopic metastatic lesions, even in the D2.0R *shCxcl10*-injected mice that effectively developed primary tumors (Supplementary Figure 2A and B). Similarly, only a single macroscopic lung metastasis was observed among the MR20 *shCxcl10*-grafted mice (Supplementary Figure 2C and D).

Due to the significant difference of primary tumor size between D2.0R *shCTL* and D2.0R *shCxcl10* tumor bearing mice (Figure 1D), which could potentially influence the development of lung metastasis analysis [38], we performed tumor cell tail vein injection experiment to directly assess their lung colonization capacity independently of the primary tumor (Figure 2A). Histopathological analysis of lungs from D2.0R *shCTL* and *shCxcl10* groups still revealed no visible macroscopic metastatic lesions (Figure 2B). Considering the possibility that disseminated cancer cells might be present as single-cells or as micrometastatic lesions below the detection threshold of histopathology, we assessed the presence of D2.0R tumor cells in the lungs by examining the expression of *Cfh, Gas6, Mme, Ogn, Tnc*, *Esr1*, previously reported as D2.0R signature genes [39]. Expression of these genes was significantly higher in the lungs of mice injected with D2.0R *shCxcl10* cells comparing with D2.0R *shCTL* cells (Figure 2C). These findings suggest that disseminated D2.0R *shCxcl10* cells can expand in the lung, albeit not to a stage of histological detection, compared to D2.0R *shCTL* cancer cells that persist at a lower burden, as disseminated, dormant cells. Strikingly, lungs from D2.0R *shCTL*-injected mice exhibited a higher density of infiltrating CD3^+^ T cells compared to those injected with D2.0R *shCxcl10* cells (Figure 2D and E), suggesting that CXCL10-mediated metastatic dormancy in the lung also correlates with a local T cell immune response--consistent with the one in the primary tumor (Figure 1J).

**Figure 2.**
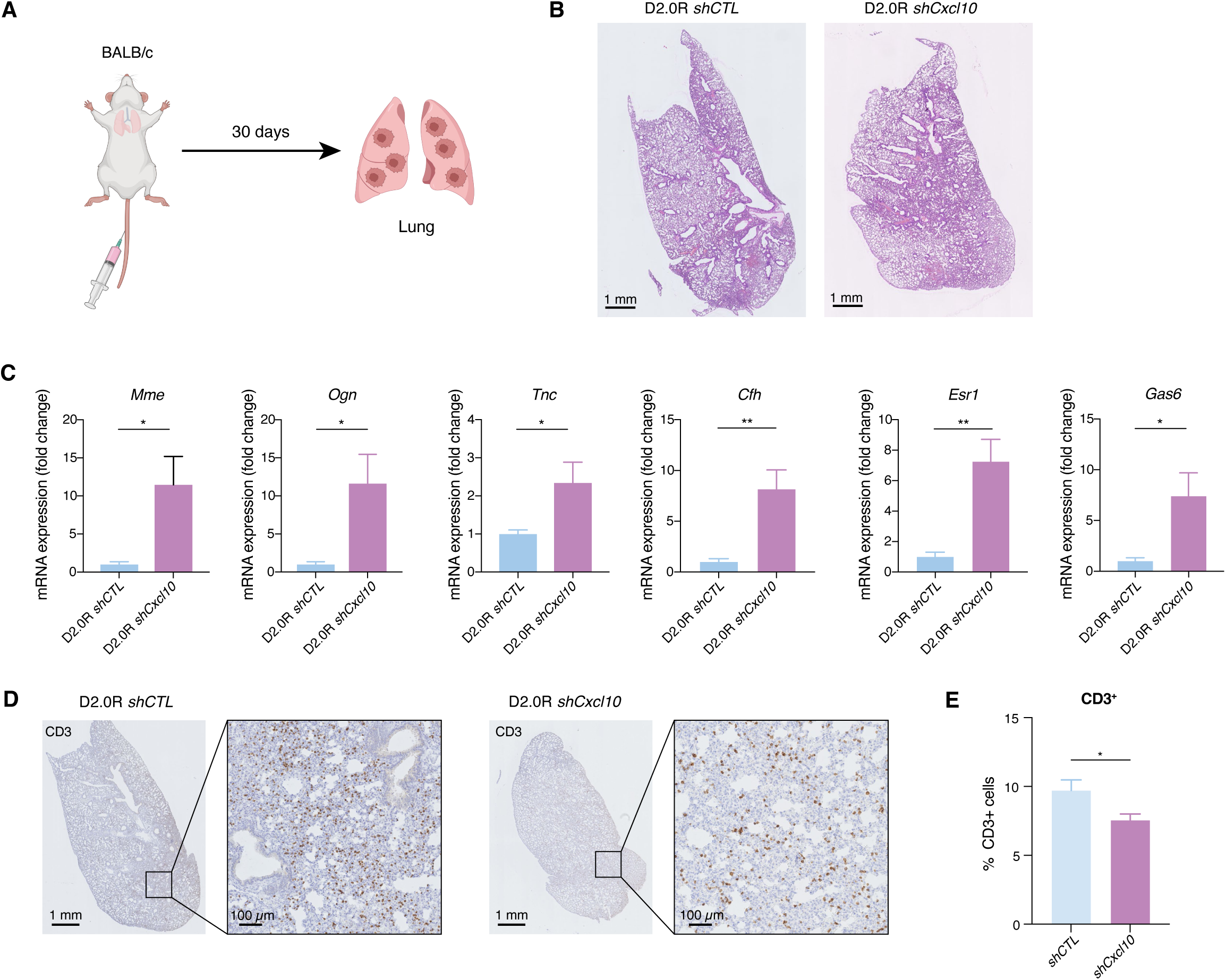
*Cxcl10* silencing enhanced lung colonization dormant tumor cells with reduced local T cell infiltration. **A** Graphical scheme showing the experimental design of the *in vivo* lung colonization assay of D2.0R-*shCTL* and D2.0R-*shCxcl10* tumor cells injected into the tail vein of BALB/c mice. **B** Representative images of H&E stained histological sections of lungs from BALB/c mice injected into the tail vein with D2.0R-*shCTL* and D2.0R-*shCxcl10* cells. n= 6/group. Scale bar: 1 mm. **C** Relative mRNA expression of D2.0R tumor signature genes, *Mme*, *Ogn*, *Tnc*, *Cfh*, *Esr1* and *Gas6,* in the lungs of D2.0R-*shCTL* and -*shCxcl10* tumor cells-injected BALB/c mice via tail vein. Expression in D2.0R-*shCT* is adjusted to 1. **D, E** Representative images of immunobiological sections (**D**) and quantification (**E**) of CD3 staining of lungs of D2.0R-*shCTL* and D2.0R-*shCxcl10* tail vein injected BALB/c mice. Scale bar: 1 mm or 100 µm. Data are represented as mean ± SEM. *P* values were calculated using unpaired two-tailed Student’s *t* test (**C & E**). ns, not significant; *, *P* < 0.05; **, *P* < 0.01.

### In vivo blockade of CXCR3 breaks tumor dormancy

CXCL10 signals via the binding to the receptor CXCR3, which is primarily expressed on activated T cells and NK cells [36, 40]. To further validate the function of the CXCL10/CXCR3 axis in immunological dormancy, we intraperitoneally (IP) injected an anti-CXCR3 blocking antibody and an IgG control antibody in D2.0R tumor bearing mice (Figure 3A-C). In anti-CXCR3 antibody-treated mice, tumor onset occurred on day 6 post-grafting and the majority of mice developing primary tumors by day 12 (Figure 3C). Only two mice in the control group developed tumors after 15 days post-grafting, while the others remained tumor-free. Analysis of immune cell populations within the tumor site showed a significant decrease in CD8^+^ T cells and a trend toward increase of MDSC, NK and DCs in the anti-CXCR3 group compared to control groups (Figure 3D). Circulating immune cell analysis revealed no significant alteration in the major immune cell populations except an increased in NK cells in anti-CXCR3 antibody-injected group compared to control group (Figure 3E).

**Figure 3.**
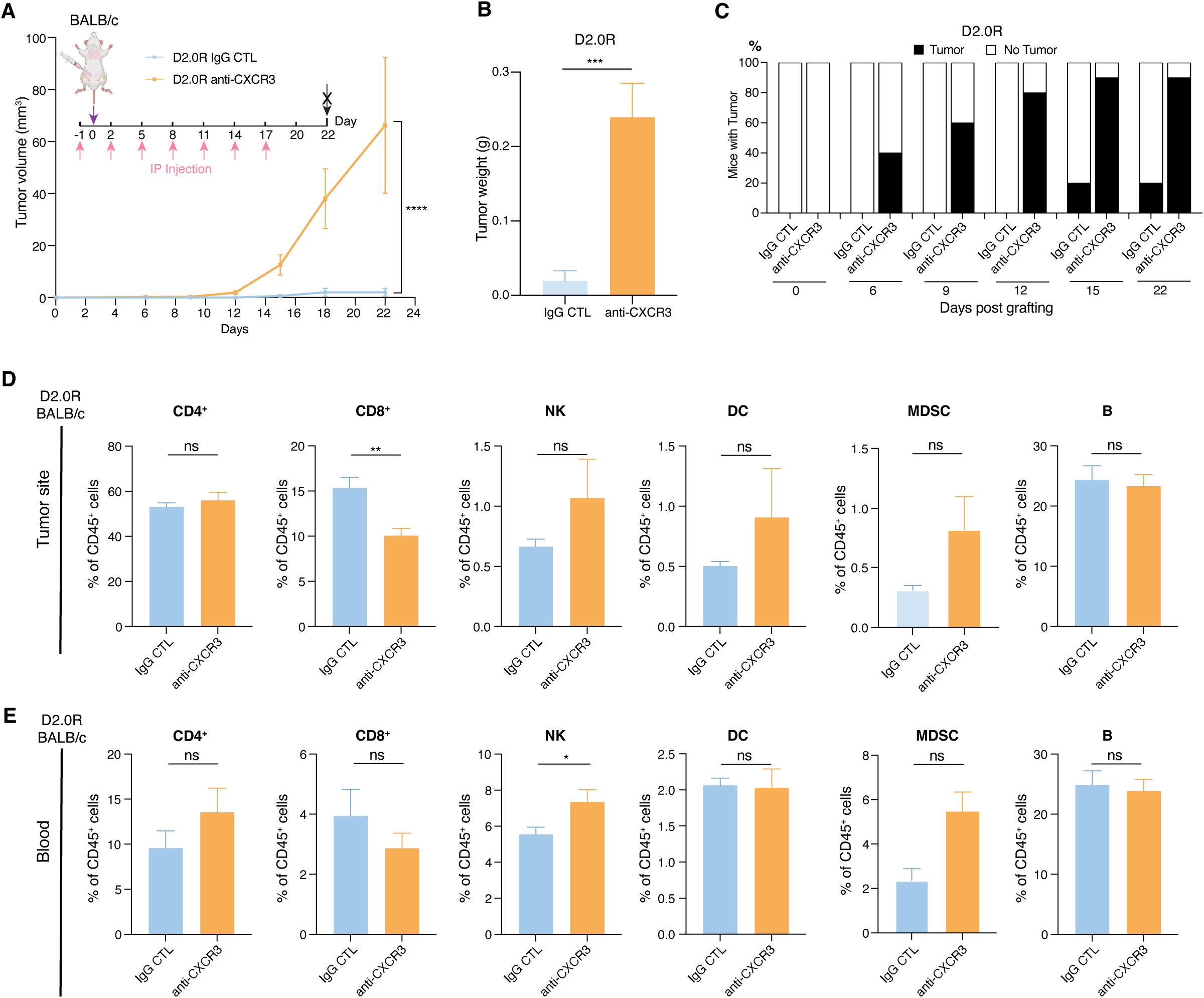
In vivo blockade of CXCR3 breaks tumor dormancy. **A-C** Primary tumor growth measured by volume (**A**), tumor weight (B) at the end of experiment and percentage of mice developing tumors over time (**C**) of D2.0R tumor cells orthotopically injected into immune competent BALB/c mice treated with neutralizing anti-CXCR3 antibodies or control IgG, as indicated. **D, E** Frequency of immune cells in the primary tumor site (**D**) or peripheral blood (**E**) detected by flow cytometry analysis of BALB/c mice orthotopically injected with D2.0R tumor cells and treated with neutralizing anti-CXCR3 antibodies or control IgG, as indicated. Data are represented as mean ± SEM. n= 9-10/group. *P* values were calculated using unpaired two-tailed Student’s *t* test (**B, D, E**) or two-way ANOVA with Sidak’s multiple-comparisons test (**A**). ns, not significant; *, *P* < 0.05; **, *P* < 0.01; ****, *P* < 0.0001.

These findings demonstrate that inhibition of CXCR3 disrupts immunological dormancy, consistent with the results obtained from D2.0R cells with *Cxcl10* silencing.

### Cxcl10 overexpression suppresses tumor growth but is not sufficient to induce dormancy

Next, we asked whether overexpression of *Cxcl10* was sufficient to induce dormancy of aggressive, non-dormant tumors. To address this question, we overexpressed *Cxcl10* in D2A1 cells (D2A1-*Cxcl10*) by lentiviral transduction (Figure 4A). Mice orthotopically injected with D2A1-*Cxcl10* cells exhibited a noticeable delay in tumor onset, albeit by the end of the experiment all mice in both groups had developed tumors (Figure 4B and C). Notably, *Cxcl10* overexpression significantly suppressed tumor growth compared to D2A1 transduced with empty vector lentivirus (D2A1-CTL) (Figure 4B and C). Flow cytometry analysis revealed a trend toward increased NK cells in D2A1-*Cxcl10* tumors compared with controls, although this difference did not reach statistical significance (Figure 4D). In contrast, more pronounced changes were observed in peripheral blood, where *Cxcl10* overexpression significantly increased NK cells and showed a trend toward higher T lymphocyte levels (Figure 4E). Meanwhile, DCs were reduced.

**Figure 4.**
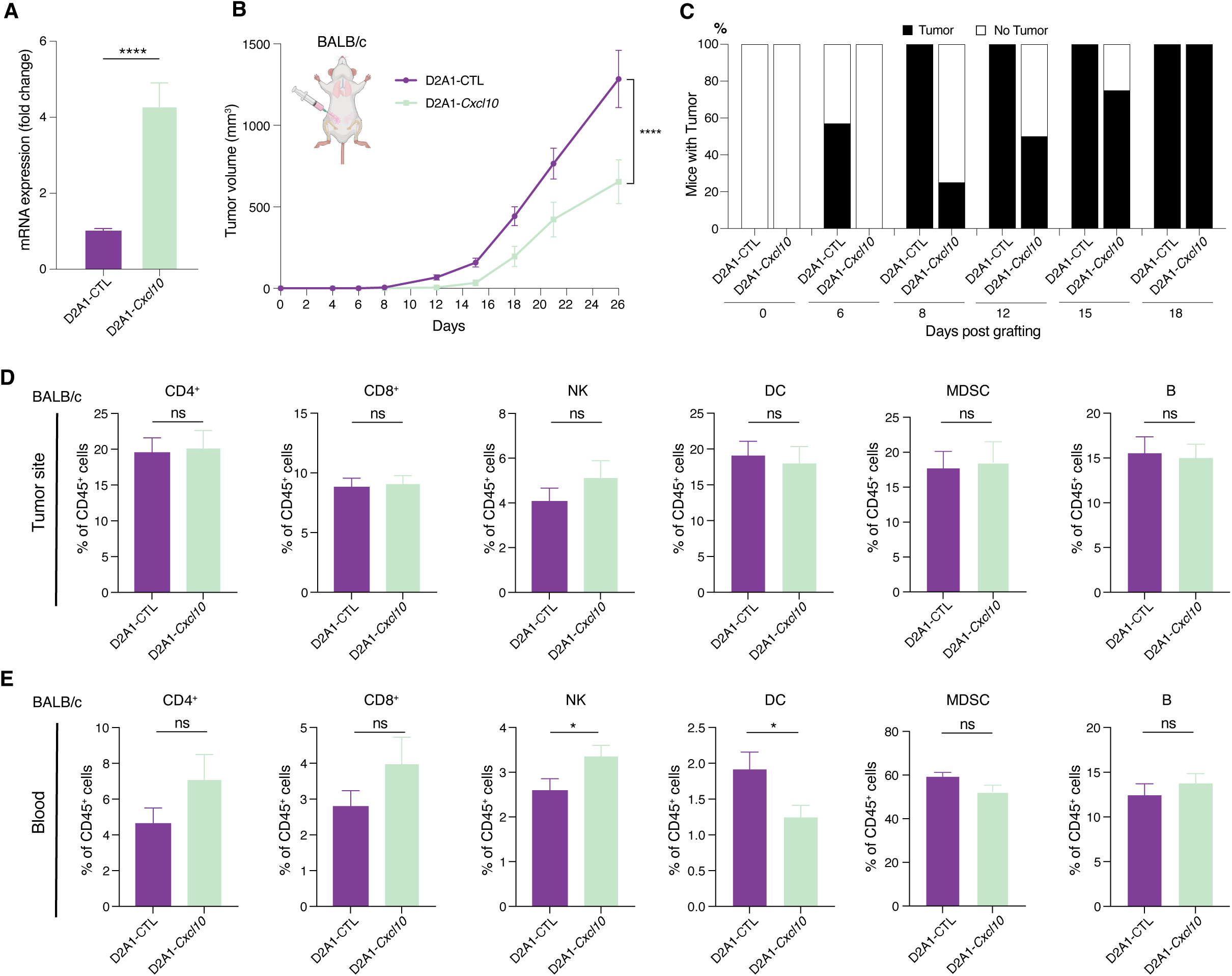
Overexpression *of Cxcl10* suppresses tumor growth. **A** Validation of lentiviral-mediated *Cxcl10* overexpression in D2A1 cells (D2A1-*Cxcl10*) by RT-qPCR. The tumor cells infected with lentivirus carrying control vector were used as control (D2A1-CTL). Expression in D2A1-CTL is adjusted to 1. **B**, **C** Primary tumor growth measured by volume (**B**) and percentage of mice developing tumors over time (**C**) in BALB/c mice orthotopically injected with D2A1-CTL or D2A1-*Cxcl10* tumor cells. n= 7-8/group. **D, E** Frequency of immune cell populations in primary tumor site (**D**) and peripheral blood (**E**). Results are expressed as percentage of the indicated cell populations within CD45^+^ cells. Data are presented as mean ± SEM. *P* values were calculated using unpaired two-tailed Student’s *t* test (**A, D, E**) or two-way ANOVA with Sidak’s multiple-comparisons test (**B**). ns, not significant; *, *P* < 0.05; ****, *P* < 0.0001.

Taken together, these results suggest that while *Cxcl10* overexpression is able to invoke anti-tumor immune response to suppress tumor growth, it is not sufficient to induce full dormancy in aggressive tumor cells.

### CXCL10-induced dormancy signature expression predicts improved survival in TNBC patients

To interrogate whether *CXCL10* expression level may affect clinic outcome in breast cancer patients, we first analyzed whether differential *Cxcl10* expression levels may have a broader impact on gene expression in cancer cells. To this end we performed transcriptomic analysis of D2.0R *shCxcl10* and D2.0R *shCTL* cells. 1293 genes were significantly differentially expressed (Supplementary Table 2). GSEA demonstrated that HALLMARK INTERFERON ALPHA and INTERFERON GAMMA pathways were the most downregulated pathways upon suppression of *Cxcl10* (Figure 5A and Supplementary Table 2). Interestingly, *Cxcl10* silencing negatively enriched for a dormancy signature extracted from the comparisons of D2.0R vs D2A1 and MR20 vs 4T1 DEG, respectively (see Methods for details) (Figure 5B and Supplementary Table 2). These data suggests that *Cxcl10* expression has a broader impact on gene expression on cancer cells, whereby high level of *Cxcl10* expression correlated with elevated interferon activity and enriched dormancy-associated gene expression.

**Figure 5.**
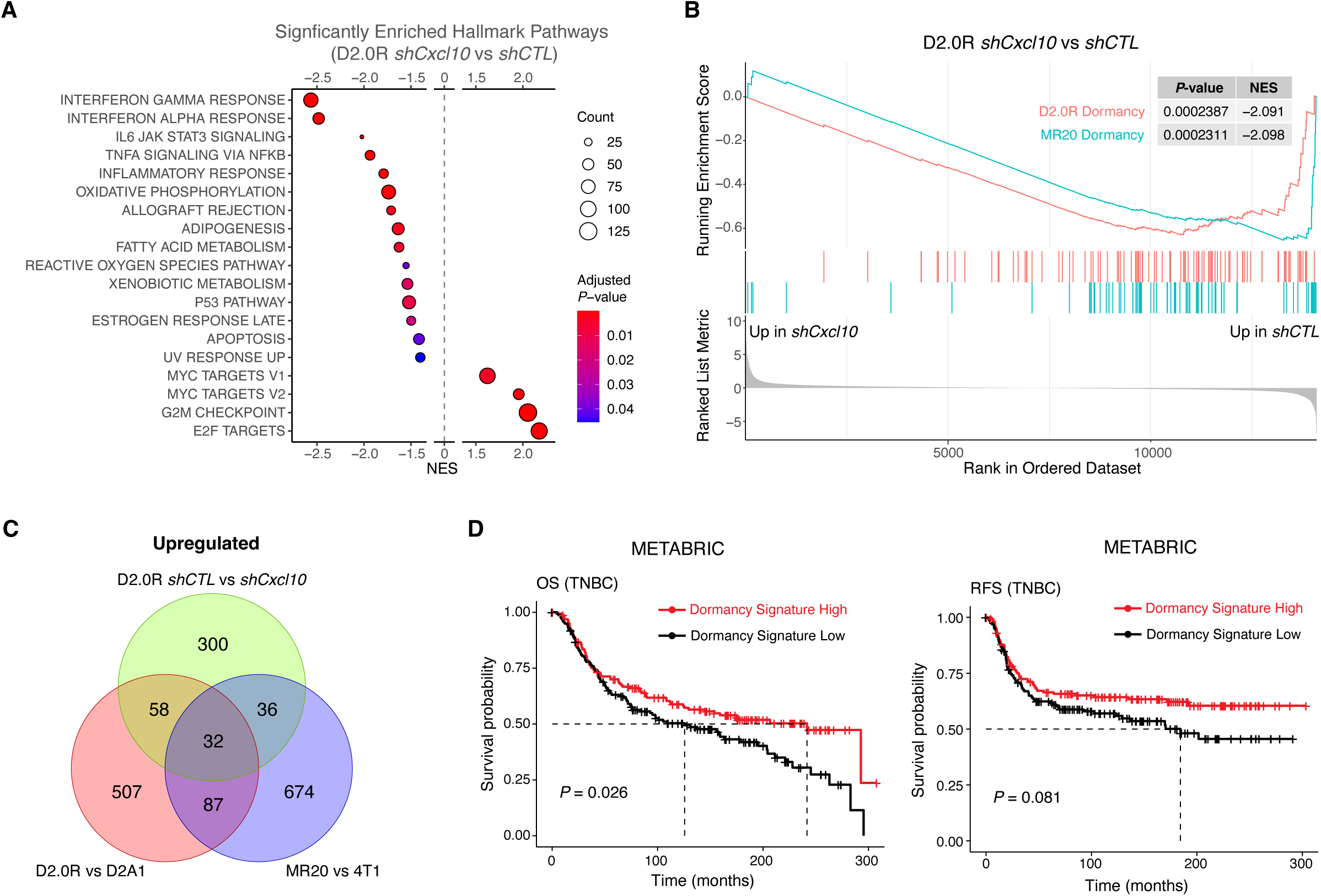
CXCL10-induced dormancy signature predicts improved survival in TNBC. **A** GSEA of RNAseq data from D2.0R-*shCxcl10* vs D2.0R-*shCTL* cells showing significantly enriched Hallmark pathways. Significance is color-coded and the number of counts indicated by the sized of the dots. **B** GSEA showing that the suppression of *Cxcl10* in D2.0R cells negatively enriched the Dormancy Signatures extracted from the comparisons of D2.0R and D2A1 (D2.0R Dormancy) and MR20 and 4T1 (MR20 Dormancy) (See details in Methods). **C** Venn diagram showing 32 common genes (Dormancy Signature) that are upregulated in dormant D2.0R and MR20 cells compared with their respective non-dormant controls, and down-regulated in D2.0R upon *Cxcl10* silencing (Up-regulated when comparing D2.0R-*shCTL* vs D2.0R-*shCxcl10*). **D** Kaplan-Meier curves showing Overall Survival (OS) and Relapse Free Survival (RFS) for TNBC patients according to high (red curve) or low (black curve) expression of the human orthologue of the Dormancy Signature in the METABRIC data sets. *P* values were calculated using log-rank test.

To validate the potential clinical significance of our findings in breast cancer dormancy, we first compared the significantly differentially expressed genes in three different datasets associated with cancer dormancy: D2.0R *vs* D2A1; D2.0R *shCTL vs* D2.0R *shCxcl10*; MR20 vs 4T1 [20] (Figure 5C and Supplementary Figure 3A). We identified 32 genes consistently upregulated in dormant tumor cells and down-regulated following *Cxcl10* silencing (upregulated in D2.0R *shCTL vs* D2.0R *shCxcl10*), defining a ‘dormancy signature’ (Supplementary Table 3). Next, we interrogated the expression of dormancy signature in METABRIC dataset comprising expression data from over 2000 breast cancer patients [41]. While the orthologous dormancy signature showed no significant association with survival in all patients (Supplementary Figure 3B), TNBC patients with higher expression of the signature exhibited significantly longer overall survival (OS; *P* = 0.0026). Intriguingly, these patients also showed a trend toward longer relapse-free survival, albeit non-significant (RFS; *P* = 0.081).

These results indicate that CXCL10-invoked immunological dormancy significantly contributes to extend survival of TNBC patients.

## Discussion

Immunological tumor dormancy represents a dynamic equilibrium where the immune system restricts tumor cell outgrowth without complete eradication, serving as a pivotal barrier against cancer progression and relapse [3]. Despite its clinical significance, the precise molecular mechanisms sustaining immunological dormancy, especially in breast cancer, are still poorly understood. Here we report that the interferon (IFN)-inducible chemokine CXCL10 is essential for sustaining immunological dormancy in TNBC.

A growing number of studies are showing that IFNs can induce tumor dormancy through multiple, mechanistically distinct pathways. Liu *et al.* demonstrated that exogenous and T cell-derived IFN-γ drives dormancy in tumor-repopulating cells through activation of the IDO-kynurenine-AhR-p27 metabolic pathway and promotion of cell cycle arrest [42]. Muller-Hermelink *et al*. reported that the combined action of T cell derived IFN-γ and tumor necrosis factor (TNF), via tumor necrosis factor p55 receptor (TNFR1), induces tumor dormancy by stimulating the release of antiangiogenic chemokines, inhibiting tumor angiogenesis and suppressing tumor cell proliferation without inducing tumor cell death [43]. More recently, Tallon de Lara and colleagues demonstrated that IFN-γ is a critical mediator of CD8^+^ T cell-induced metastatic dormancy in breast cancer [13].

We have previously reported that the sustained, autocrine activation of the IRF7/IFN-β/IFNAR axis in cancer cells induced a state of immunological dormancy in a model of spontaneous breast cancer dormancy (D2.0R) and a model of chemotherapy induce breast cancer dormancy (MR20). Activation of the IRF7/IFN-β/IFNAR axis promoted the infiltration and activations of CD4^+^ and CD8^+^ T cells in the tumor microenvironment, while inhibiting the recruitment of MDSCs. Conversely, inhibiting the IRF7/IFN-β/IFNAR axis in tumor cells reversed tumor dormancy. Depletion of CD4^+^ or CD8^+^ T lymphocytes prevented dormancy confirming the key role of the immune system [20]. This observation raised the question of the mechanism by which IFN induced immunological dormancy in these models.

Here we addressed this question by analyzing the transcriptome of dormant tumor cells (D2.0R and MR20) compared to their non-dormant counterparts (D2A1 and 4T1). Consistently, we observed that the IFN-induced chemokines *Cxcl10*, *Cxcl10*, and *Cxcl11* were significantly upregulated in dormant tumor cells. Among these, *Cxcl10*, also known as Interferon induced Protein 10 (IP10), was the one most predominantly expressed. Targeting *Cxcl10*, either by genetic silencing in cancer cells, or systemic antibody-mediated inhibition of its receptor CXCR3, reactivated dormant tumor cells *in vivo*, resulting in clinical tumor growth. Importantly, *Cxcl10* silencing decreased tumor-infiltration by CD4^+^ or CD8^+^ T lymphocyte and increased tumor-infiltration by MDSC, consistent with immunological dormancy. Furthermore, dormant cells effectively formed tumors in immune-deficient mice independently of *Cxcl10* status, demonstrating that the CXCL10/CXCR3 axis-mediated dormancy requires a functional immune system

CXCR3 chemokine ligands CXCL9 and CXCL10 are pivotal mediators of anti-tumor immunity by critically orchestrating the recruitment and activation of effector immune cells including CD8^+^ T cells within the tumor microenvironment [44, 45]. In a study using murine models of head and neck squamous cell carcinoma (HNSCC), Mikucki *et al.* demonstrated that activation of the CXCR3 receptor by its ligand CXCL10 significantly enhances the trafficking of T cells into the tumor microenvironment and their activation, resulting in more effective tumor cell killing [46]. Hoch *et al.* by mapping the spatial distribution of chemokines and immune cells within patient metastatic melanoma tissues, found that areas rich in CXCL9 and CXCL10 chemokines closely associate with clusters of exhausted CD8^+^ T cells feeding a forward loop where newly recruited T cells produce IFN-γ, stimulating further *CXCL9* and *CXCL10* production and attracting more immune cells [47]. One study using an *ex vivo* model of liver metastasis, however, reported that CXCL10 can trigger the emergence of dormant TNBC cells within the liver, possibly *via* activation of liver cells [48].

Recent evidence indicates that the CXCL10/CXCR3 axis is crucial in modulating the efficacy of immunotherapy [25, 49, 50]. Peng *et al.* showed that blocking PD-1 in a murine melanoma model enhances adoptive cell transfer therapy by increasing IFN-γ expression at the tumor site resulting in *CXCL10* production promoting T-cell infiltration and improving antitumor response [27]. Ayers and colleagues showed that an IFN-γ-related gene signatures, including *CXCL9* and *CXCL10,* can predict tumor responsiveness to PD-1 checkpoint blockade in several cancer types including breast cancer [51]. Chow *et al.* demonstrated that CXCR3 plays a critical role in promoting intratumoral CD8^+^ T cell proliferation and function following anti-PD-1 immunotherapy in melanoma, and *CXCL9* and *CXCL10* expression correlated with improved treatment outcome [52]. Additionally, *IFNs* and *CXCL10* expression were also shown to be correlated with better clinical outcomes after chemotherapy in breast cancer [53].

Here we provide the first compelling evidence that the CXCL10/CXCR3 axis also play a critical role in maintain breast cancer dormancy. Disruption of CXCL10/CXCR3 axis reduced CD4^+^ and CD8^+^ T cell infiltration, increased MDSC infiltration in the primary tumor, awakened dormant cells to form primary tumors, and increased the burden of disseminated tumor cells (DTC) in the lungs. Notably, while *Cxcl10* silencing induced dormancy escape at the primary site, only DTC were detected in lung tissue, without histological evidence of macrometastases formation. The limited progression of DTC in the lung may be due to the short time frame of the experiment or likely the requirement of additional factors for full metastatic outgrowth, in particular the formation of a supportive metastatic niche [54, 55]. Nevertheless, this effect was accompanied by reduced T cell infiltration in the lungs, suggesting that *Cxcl10* silencing does suppress immune surveillance in the lung, alike in the primary tumor site.

These observations raised the question of whether increasing expression of *Cxcl10* in aggressive cancer cells was sufficient to induce dormancy. We addressed this question experimentally by overexpressing *Cxcl10* in aggressive D2A1 cells. While *Cxcl10* overexpression significantly blunted primary tumor growth, it failed to induce complete dormancy. This outcome is consistent with the modest alterations observed in both local and systemic immune responses. Additional cooperating mechanisms, beyond *Cxcl10*, are likely required to establish a durable dormant state. Further studies are needed to fully dissect the complexity of immunological tumor dormancy before translating our findings into a therapeutic intervention to induce or maintain dormancy in patients at high risk for progression.

The identification of a *Cxcl10*-induced dormancy-associated signature has nevertheless a relevant clinical implication: patients with a high dormancy signature have a better OS compared to patients with a low dormancy signature who have a worse OS. Importantly, the difference in OS appears starting 3 years post diagnosis, consistently with a dormancy effect. Strikingly, the dormancy signature is only significant in TNBC, bringing further support to the growing evidence that dormancy is not limited to ER^+^ breast cancer but may also occur in TNBC [20, 54, 56–59]. This implies that TNBC patients with a low dormancy signature should be closely monitored also late after initial therapy, alike patients with ER^+^ breast cancer [60, 61].

In conclusion, our study provides original experimental evidence that the CXCL10/CXCR3 axis is essential for sustaining immunological tumor dormancy in TNBC by modulating the immune tumor microenvironment. The identification of a CXCL10-dependet dormancy signature that stratify TNBC patients for better OS late after initial therapy has direct clinical implication calling for a closer monitoring late after initial therapy of TNBC patients with a low signature.

## Methods

### Cell culture

The murine breast cancer cell lines D2A1 and D2.0R were generously supplied by Dr Jonathan Sleeman (Medical Faculty at Heidelberg University, Mannheim, Germany). These cell lines were maintained in high glucose Dulbecco’s Modifier Eagle Medium (DMEM) supplemented with 10% heat-inactivated fetal bovine serum (FBS) and 1% penicillin-streptomycin (P/S, Life Technologies – Invitrogen) and 1% Non-essential Amino Acid (NEAA) (Gibco). The 4T1 mouse mammary carcinoma cell line, kindly provided by Dr Fred R. Miller (Michigan Cancer Foundation, Detroit, MI, USA) was also cultured in the same medium. MR20 cell line, derived in-house [20] was cultured in high glucose DMEM supplemented with 10% heat-inactivated FBS, 1% P/S, with Methotrexate added at 20 ng/ml every 2 to 3 days to maintain selective pressure.

### Lentiviral constructs and gene modulation by shRNA and cDNA

Short hairpin RNA (shRNA) and cDNA sequences targeting CXCL10 (*shCxcl10*) and non-targeting sequence (*shNT*) in pLKO.1-puro lentiviral constructs were purchased from Nucleus Biotech. Lentivirus production was performed using a standard calcium phosphate transfection method in HEK 293T cells. Cells were cultured in DMEM and co-transfected with three plasmids: the transfer vector carrying the shRNA/cDNA of interest, the packaging plasmid pSD16, and the envelope plasmid pSD11. The total DNA was combined with 0.5 M CaCl₂ and 2X HeBS buffer. The DNA mixture was incubated for 10 minutes to allow precipitate formation before adding it to the HEK 293T cells. After overnight incubation, the transfection medium was replaced with fresh complete medium (6–8 mL), and viral supernatants were collected 48–72 hours post-transfection. Viral particles were either used fresh or stored at −80 °C for later use. Transduction of target cells was performed by incubating them with viral supernatant in the presence of polybrene (8 µg/mL) to enhance infection efficiency. Cells were selected for stable integration of viral constructs by antibiotic resistance with puromycin. Gene knockdown or overexpression efficiency was validated by Real-time reverse transcription qPCR (qRT-PCR).

### Tumor models

For orthotopic models, D2A1, D2.0R and MR20 cells (5 x 10^4^ cells in 50 µl PBS/10% of 8.1 mg/ml Matrigel mixture per mouse) were injected in the fourth right mammary glands of 6–7-week-old BALB/c (Invitrogen), and NOD SCID common gamma 2 chain deficient (NSG, University of Lausanne, Switzerland) female mice. Tumor growth was measured three times a week with caliper and tumor volume was calculated by the equation: volume = (length × width^2^) × π/6. At the endpoint, mice were sacrificed according to defined ethical criteria with lethal anesthesia for terminal blood, lung and tumor collection. For pharmacologic inhibition experiment intraperitoneal injection of anti-CXCR3 antibody (200 µg/mouse; clone CXCR3-175; BioXcell) and isotope control antibody (100 µg/mouse; #BE0091; BioXcell) began one day prior to orthotopic grafting of D2.0R WT cells intro the mammary fat pad of BALB/c female mice and continued every three days for a total of seven injections. For tail vein model, 2 x 10^5^ D2.0R *shCTL* and D2.0R *shCxcl10* cells were injected in 50 µl of PBS. All animal procedures were performed in accordance with the Swiss legislation on animal experimentation and approved by the Cantonal Veterinary Service of the Canton Fribourg (2021-29_FR).

### Histopathology

At the end of the *in vivo* experimental procedures, both tumor and lung tissues were collected, fixed in formalin, and subsequently embedded in paraffin. Serial sections with a thickness of 5 μm were prepared from the embedded tissue blocks. For analysis, 3 to 4 sections spaced 100 µm apart were selected and stained using hematoxylin and eosin (H&E) to evaluate tumor/tissue structure and to measure the extent of lung metastases. The tissue sections were digitized using a Nanozoomer scanner (Hamamatsu Photonics), and metastatic sites were manually enumerated using the NDP.view2 software (Hamamatsu Photonics). For immunostaining, antigen retrieval was performed by boiling the tissue sections in appropriate buffer for 20 minutes. Endogenous peroxidases were blocked by incubation with 0.6% H_2_O_2_ in methanol. After blocking with PBS-10% BSA, sections were incubated with anti-CD3 MAb (ab5690, Abcam) overnight at 4°C and, after a washing step, with the Goat-anti-Mouse-HRP conjugate secondary antibody for 2 hours at room temperature. Dako Envision^+^ was used with diaminobenzidine (DAB) tablets (Sigma-Aldrich) to detect HRP. Images were taken with a widefield microscope (Leica).

### Flow cytometry analysis

For solid tissues, both tumor and lung tissues were dissected and processed immediately. Enzymatic digestion was performed on previously cut small pieces of tumor and lungs, using serum-free medium containing DNase and collagenase I (Roche), then incubated at 37°C for 45 minutes on a shaking platform. The tissue suspensions were then filtered through 70 µm sterile nylon gauzes and the resulting cell suspensions were centrifuged at 400 g for 5 minutes. For blood analysis, peripheral blood samples were collected from lethally anesthetized mice, using standard EDTA anti-coagulation procedures to prevent clotting. Red blood cells were lysed using ACK lysis buffer and leukocyte pellets were recovered after centrifugation at 1400 rpm for 5 minutes. Cell pellets from blood, lungs and tumor were resuspended in PBS and counted with automated cell counter LUNA-II™ (Logos Biosystems). Two million cells were stained with viability dye for 20 min at 4°C to exclude dead cells from analysis. Stained cells were washed and centrifuged to remove excess viability dye before staining with a panel of fluorophore-conjugated antibodies targeting key immune cell markers (listed below). Stained cells underwent additional washing steps and centrifugation to remove excess antibodies and resuspended in FACS buffer before acquisition with the flow cytometer Cytek Aurora (Cytek Biosciences). Data were analyzed by FlowJo, version 10.0.7 (Tree Star Inc.).

### Flow cytometry antibodies

The following anti-mouse antibodies were used according to the manufacturer’s instructions: anti-CD16/CD32 Fc blocking antibody (BD Bioscience), CD45R/B220-BV785 (clone RA3-6B2), Ly-6G-Alexa Fluor 647 (clone 1A8), CD11b-BV650 (clone M1/70), CD11c-Alexa Fluor 488 (clone N418), CD4-PE (clone GK1.5), CD49b-Pacific Blue (clone DX5), Ly-6G/Ly-6C-APC/Cy7 (clone RB6-8C5), CD8a-PE-Cy7 (clone 53-6.7), CD3-BV570 (clone 17A2, Biolegend), CD45-BUV737 (clone 30-F11, BD Bioscience), Live/Dead fixable blue dead cell stain kit (Thermofisher).

### Real-time reverse transcription qPCR and primers

Analysis of mRNA expression levels was performed using real-time PCR. Total RNA was extracted from adherent cells using the RNA Plus extraction kit (Machery-Nagel), following the manufacturer’s protocol. For each sample, 1 µg of RNA was reverse-transcribed into cDNA using the SuperScript II Reverse Transcriptase Kit (Life Technologies – Invitrogen) according to the manufacturer’s instructions. The qPCR reactions were carried out using SYBR® Master Mix (SensiFAST SYBR HI-ROX Kit, Meridian bioscience) on a StepOnePlus™ Real-Time PCR System (Applied Biosystems, Life Technologies – Invitrogen). Gene expression levels were normalized to the murine GAPDH housekeeping gene, and relative quantification was calculated using the comparative Ct (ΔΔCt) method. The following murine-specific primers were purchased from Microsynth AG: *GAPDH* forward, 5’-CATCACTGCCACCCAGAAGACTG-3’, reverse 5’-ATGCCAGTGAGCTTCCCGTTCAG-3’; CXCL9 forward, 5’-CCTAGTGAT AAGGAATGCACGATG-3’, reverse 5’-CTAGGCAGGTTTGATCTCCGTTC-3’; *CXCL10* forward, 5’-ATC ATCCCTGCGAGCCTATCCT-3’, reverse 5’-GACCTTTTTTGGCTAAACGCTTTC-3’; *CXCL11* forward, 5’-CCGAGTAACGGCTGCGACAAAG-3’, reverse 5’-CCTGCATTATGAGGCGAGCTTG-3’; *Cfh* forward, 5’-GAGACAAGCAGGAGTACGAACG-3’, reverse 5’-CCATCCAAGTATTTCACGGTGGT-3’; *Gas6* forward 5’-GAACTTGCCAGGCTCCTACTCT-3’, reverse 5’-GGAGTTGACACAGGTCTGCTCA-3’; *Mme* forward 5’-CAGCCGAAACTACAAGGAGTCC-3’, reverse 5’-CATAAAGCCTCCCCACAGCATTC-3’; *Ogn* forward 5’-AACGACCTGGAATCTGTGCCTC-3’, reverse 5’-TCGCTCCCGAATGTAACGAGTG-3’; *TNC* forward 5’-GAGACCTGACACGGAGTATGAG-3’, reverse 5’-CTCCAAGGTGATGCTGTTGTCTG-3’; *Esr1* forward 5’-TCTGCCAAGGAGACTCGCTACT-3’, reverse 5’-GGTGCATTGGTTTGTAGCTGGAC-3’.

### RNA sequencing and data analysis

Three or four independent batches of D2A1, D2.0R, D2.0R-shCTL and D2.0R-shCXCL10 cells were used for RNA extraction using the NucleoSpin RNA extraction kit (Macherey-Nagel) following manufacture’s manual instructions. Samples were normalized for 1 μg RNA in a volume of 20 μl, sequenced on the NextSeq500 sequencer using the NextSeq 500/550 HT reagent v2 kit (Illumina) at the Lausanne Genomics Technologies Facility (GTF, UNIL) in Lausanne, Switzerland. For data analysis, all sequencing reads were processed for quality control, removal of low-quality reads, adaptor sequence and ribosomal RNA by RSeQC (v5.0.1) [62], Cutadapt (v4.8) [63] and Fastq Screen (v0.11.3) [64]. The filtered reads were aligned against Mus_musculus GRCm39.110 genome using STAR (v2.7.10a) [65] and summarized with htseq-count (v0.11.2) [66] using gene annotation. The normalization of the read counts and the analysis of the differential expression between the groups of samples were performed with in R (v4.4.3), a free software environment available at https://www.r-project.org/ using packages DESeq2 (v1.48) [67].

GSEA pathway enrichment analysis was performed using packages clusterProfiler (v4.16.0) [68] with the input of fold change-ranked gene list from DESeq2 comparing the Hallmark genesets from MSigDB (v2024.1) [69] or customized D2.0R dormancy and MR20 dormancy signature with default settings. The significant altered pathways were determined by computing moderated t-statistics and false discovery rates with the limma (v3.64.1) [70] for pairwise comparison. The plots were produced with R packages ggplot2 (v2.5.2) [71] or enrichplot (v1.28.4) [72].

D2.0R dormancy signature was extracted from the top 100 most up-regulated genes by comparing D2.0R vs D2A1 with the threshold of adjusted P value < 0.05, fold change > 2 and average normalized count number > 100. MR20 dormancy signature was extracted from top 100 most up-regulated genes by comparing MR20 vs 4T1 with threshold of a of adjusted P value < 0.05 and fold change > 2 by analyzing publicly available dataset GSE100973 [20].

For Venn diagram, the genes fulfilling the threshold of adjusted *P* value < 0.05, fold change > 1.5 or < -1.5 and average normalized count number > 50 for RNAseq data are compared. The figures were produced with R package venn (v1.12) [73].

### Clinical data analysis

To validate our finding with human data, the human orthologs of murine dormancy signature genes were used. Conversion from murine to human gene symbols was performed with the biomaRt package (v2.64.0) [74, 75], using the reference mart https://dec2021.archive.ensembl.org. Zscore normalization and signature assessment was performed using the hacksig package (v0.1.2) [76]. Molecular Taxonomy of Breast Cancer International Consortium (METABRIC) breast cancer data was downloaded from cBioPortal [77, 78] in August 2024. The expression values were stratified in two groups by median values. The tumor subtype was characterized based on the status of ER, PR and HER2 included in the metadata, and 320 patients were classified as TNBC. Survival curves were generated using the ggsurvplot function from the survminer package (v0.5.0) [79], and were compared between groups using Cox proportional hazard regression model performed through the coxph and survfit functions from the survival package (v3.8-3) [80].

### Statistical analysis

All statistical analyses were conducted using GraphPad Prism software version 7.0. Data are presented as mean ± standard error of the mean (SEM) from at least three independent experiments, as indicated in figure legends. For comparisons between two groups, either an unpaired Student’s *t* test or the Mann–Whitney U test was employed. For comparisons involving more than two groups, two-way analysis of variance (ANOVA) followed by Sidak’s multiple comparison were used. A *P* value less than 0.05 was considered statistically significant. RNAseq data were presented as mean of normalized count numbers ± SD and Wald test included in R package DESeq2 was applied.

## Data availability

The transcriptomic data generated by this study have been deposited in the GEO database under the access code GSEXXXXXX. The previous published dataset GSE100973 were downloaded from GEO and re-analyzed as described in the Methods.

## Acknowledgments

This work was supported by a grant of the Swiss National Science Foundation to C.R. (310030_208136). The authors wish to thank Sarah Cattin for assistance with FACS analysis, Jean-Christophe Stehle and Janine Horlbeck for tissue staining, and Félix Meyenhofer for assistance with microscope. Some elements in the illustrative figures are created in BioRender. Lan, Q. (2026) https://BioRender.com/gka0rc9.

---

**Supplementary Figure 1.**
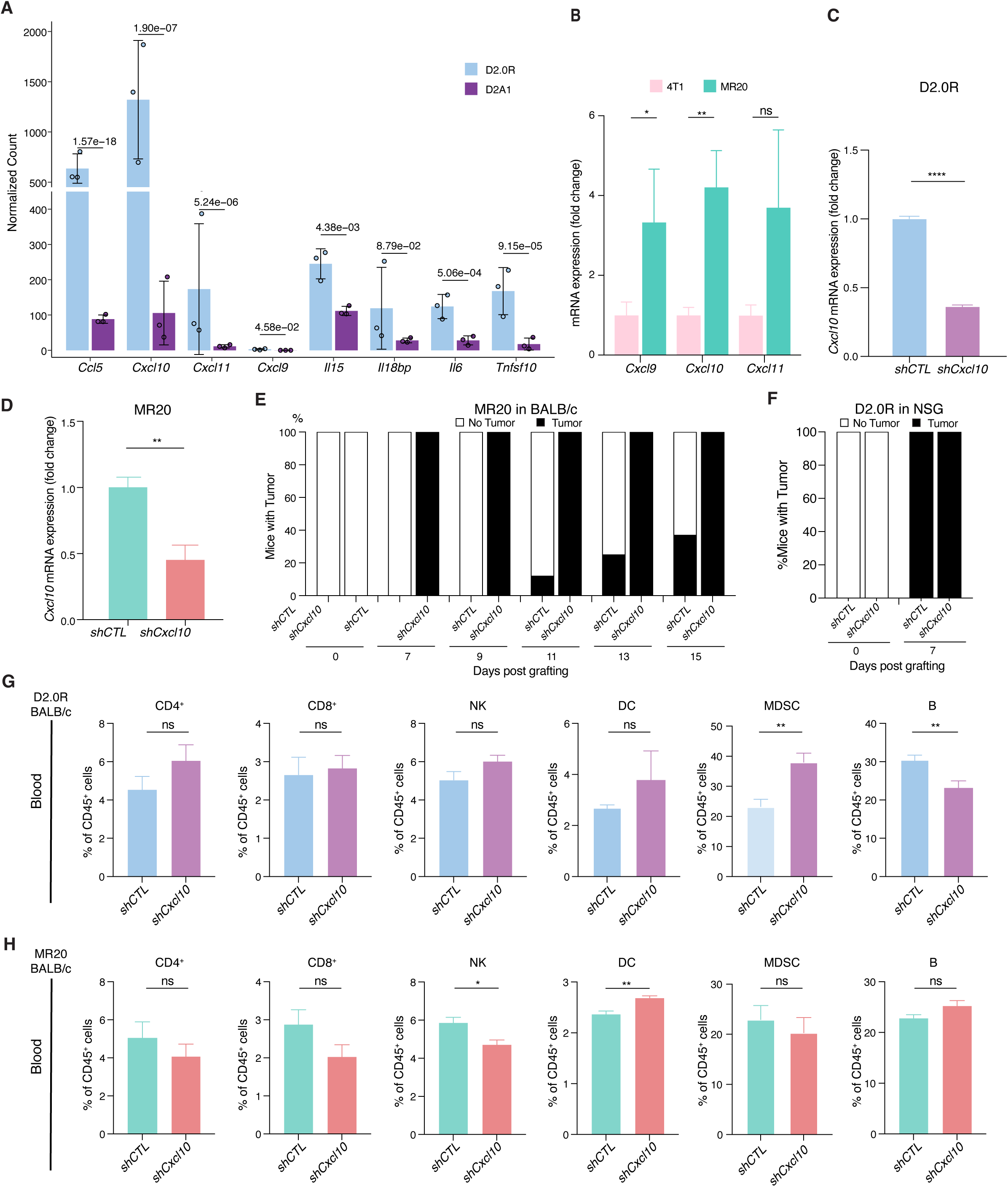
Selected gene expression analysis in D2A1 vs D2.0R and effect of Cxcl10 silencing D2.0R cells on tumor incidence and local immune cell environment. **A** Normalized RNA-seq counts of the indicated ligand coding genes that belong to the INTERFERON ALPHA and GAMMA RESPONSE pathways and are differentially expressed between D2.0R and D2A1 cells. **B** Relative mRNA expression of *Cxcl9*, *Cxcl10* and *Cxcl11* in MR20 and 4T1 cell lines examined by RT-qPCR. n= 2-3/group. **C, D** Validation of lentiviral-mediated *Cxcl10* expression silencing in D2.0R cells (**C**) and MR20 (**D**) measured by by RT-qPCR. The tumor cells infected with lentivirus carrying non-targeting control vector were used as control (*shCTL*) and expression level set to 1. n = 3-4/group **E** Percentage of immune-competent BALB/c mice developing tumors over time following orthotopic implantation of *shCTL*- or *shCxcl10*-transduced MR20 tumor cells. n= 7-8/group. **F** Percentage of immune-deficient NSG mice developing tumors over time following orthotopic implantation of *shCTL*- or *shCxcl10*-transduced D2.0R tumor cells. n= 7-8/group. **G, H** Fraction of immune cell populations in peripheral blood of BALB/c mice orthotopically injected with *shCTL* or *shCxcl10* tumor cells derived from D2.0R (**G**) or MR20 (**H**), as determined by flow cytometry analysis. Results are expressed as percentage of the indicated cell populations within CD45^+^ cells. n= 6-8/group. For panel **A**, the data are presented as mean ± SD, and *P* value are calculated using Wald test. The data in **B**-**D**, **G** and **H** are represented as mean ± SEM, and *P* values were calculated unpaired two-tailed Student’s *t* test. ns, not significant; *, *P* < 0.05; **, *P* < 0.01; ****, *P* < 0.0001.

**Supplementary Figure 2.**
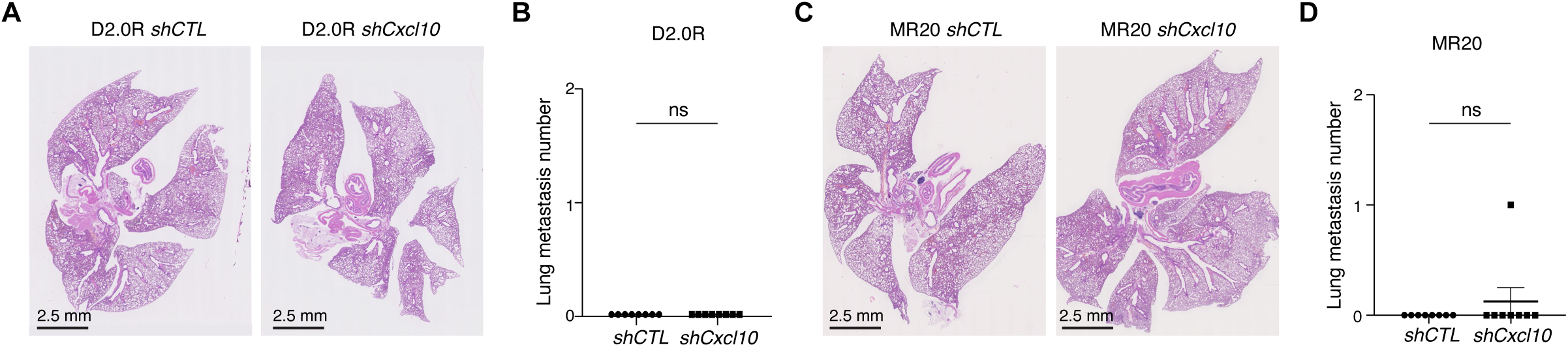
Suppression of *Cxcl10* has limited effect on lung metastasis. Representative histological images of lung (H&E staining) and corresponding quantifications of metastatic burden from BALB/c mice orthotopically injected with D2.0R*-shCTL* or D2.0R*-shCxcl10* tumor cells (**A** & **B**) and MR20*-shCTL* or MR20-*shCxcl10* tumor cells (**C** & **D**). Scale bar: 2.5 mm. n = 8/group. Data are represented as mean ± SEM. *P* values were calculated using Mann-Whitney test. ns, not significant.

**Supplementary Figure 3.**
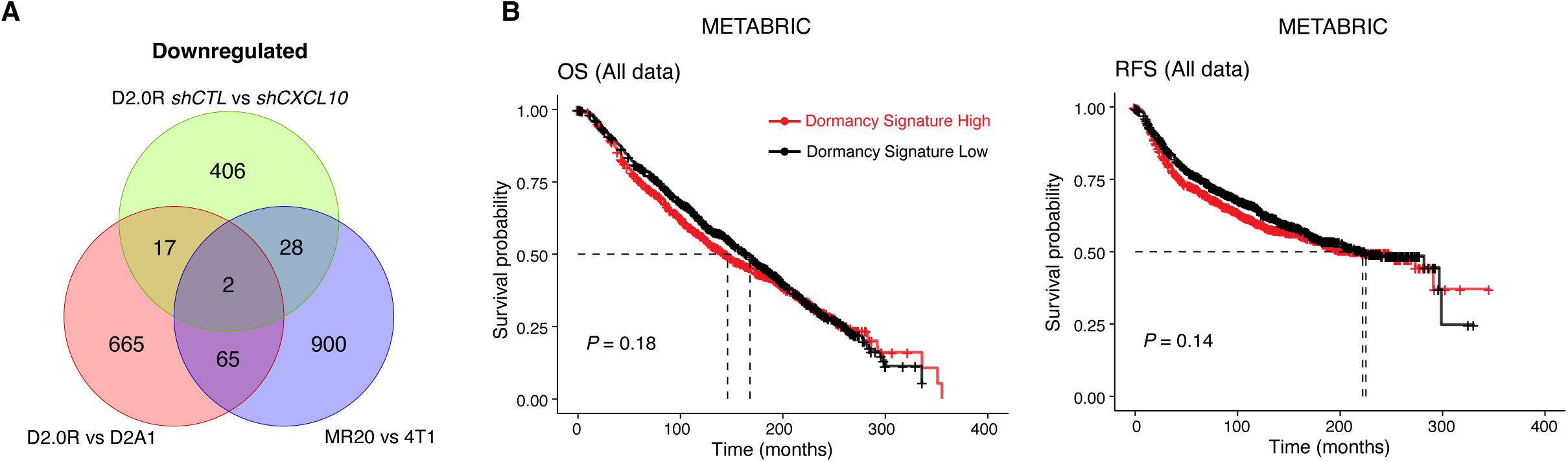
CXCL10-induced dormancy signature expression in all breast cancer patients and effect on OS and RFS. **A** Venn diagram showing common genes that are downregulated in dormant D2.0R and MR20 cells compared with their respective non-dormant controls and upregulated in D2.0R upon *Cxcl10* knocking down (downregulated when comparing D2.0R *shCTL* vs *shCxcl10*). **B** Kaplan-Meier curves showing OS and RFS for all patients according to high or low expression of human orthologue of Dormancy Signature in METABRIC data sets. *P* values were calculated using log-rank test.

